# A Continuous Extension of Gene Set Enrichment Analysis using the Likelihood Ratio Test Statistics Identifies VEGF as a Candidate Pathway for Alzheimer’s disease

**DOI:** 10.1101/2023.08.22.554319

**Authors:** Ali Mahzarnia, Michael W. Lutz, Alexandra Badea

**Affiliations:** Department of Radiology, Duke University School of Medicine, Durham, NC, USA; Department of Neurology, Duke University School of Medicine, Durham, NC, USA; Biomedical Engineering, Duke University, Durham, NC, USA; Brain Imaging and Analysis Center, Duke University School of Medicine, Durham, NC, USA

**Keywords:** Continuous GSLRT, Likelihood Ratio-Derived Test Statistics, Alzheimer’s Disease, RNA-Seq

## Abstract

**Background:** Alzheimer’s disease involves brain pathologies such as amyloid plaque depositions and hyperphosphorylated tau tangles and is accompanied by cognitive decline. Identifying the biological mechanisms underlying disease onset and progression based on quantifiable phenotypes will help understand the disease etiology and devise therapies.

**Objective:** Our objective was to identify molecular pathways associated with AD biomarkers (Amyloid-β and tau) and cognitive status (MMSE) accounting for variables such as age, sex, education, and APOE genotype.

**Methods:** We introduce a novel pathway-based statistical approach, extending the gene set likelihood ratio test to continuous phenotypes. We first analyzed independently each of the three phenotypes (Amyloid-β, tau, cognition), using continuous gene set likelihood ratio tests to account for covariates, including age, sex, education, and APOE genotype. The analysis involved a large sample size with data available for all three phenotypes, allowing for the identification of common pathways.

**Results:** We identified 14 pathways significantly associated with Amyloid-β, 5 associated with tau, and 174 associated with MMSE. Surprisingly, the MMSE outcome showed a larger number of significant pathways compared to biomarkers. A single pathway, vascular endothelial growth factor receptor binding (VEGF-RB), exhibited significant associations with all three phenotypes.

**Conclusions:** The study’s findings highlight the importance of the VEGF signaling pathway in aging in AD. The complex interactions within the VEGF signaling family offer valuable insights for future therapeutic interventions.

## Introduction

One strategy to better understand and accurately model late-onset Alzheimer’s disease, is to incorporate various genetic, clinical, and environmental factors into a cohesive model. This model should establish connections between measurable biomarkers and risk factors. Several key factors play a significant role in shaping the risk of Alzheimer’s disease (AD). These factors encompass both genetic influences, such as APOE genotype and sex, as well as environmental elements, including education level, diet, and age. In recent years, the identification and characterization of AD have been facilitated by the use of biomarkers like Amyloid-β (Aβ), phosphorylated tau (tau), and measures of neurodegeneration. Furthermore, AD is distinguished by memory impairment, often evaluated using the Minimental State Evaluation (MMSE).

AD has a deleterious impact on American lives, as over 6 million individuals are currently afflicted with Alzheimer’s, with a projected twofold increase by 2050. The disease’s high mortality rate claims 1 in 3 seniors while imposing significant economic burdens, costing the nation billions [1]. However, the biological background conducive to developing AD remains unknown. The aim of this study was to address the existing knowledge gap by pinpointing molecular pathways that play a crucial role in modulating levels of hallmark AD pathologies, as well as memory function.

Recent publications have revealed a role for multiple pathways in AD, based on brain proteomics and transcriptomic analyses [2, 3]. These studies have revealed pathways that are relevant not only to neurons but also to cells regulating response to inflammation [4], such as microglia [5]. Interestingly endothelial cells, astrocytes and neurons that control neurovascular functions have been shown to play an important role in AD [6]. Other cell types and subcellular components such as mitochondria may be involved [7]. [8] identifies a novel brain-enriched RING finger E3 ligase, RNF182, which shows elevated expression in AD brains and may play a role in controlling neurotransmitter release. Pathways involved with filament-based processes, cellular detoxification, and wound healing have also been involved [2, 3]. A decline in sensory function, including taste has also been reported with aging and AD [9]. Importantly, the vascular endothelial growth factor has been associated with AD (VEGF) [10, 11], and while its role in neurodegeneration is yet to be understood, it presents a druggable target for therapies. However, most studies focused on comparisons of two or three groups of subjects using discrete classification variables, such as case/control, without accounting for the relationships between multiple hallmark biomarkers. Here we propose an approach to detect common gene pathways based on RNA-Seq changes associated with continuous-scale changes in multiple biomarkers and clinical phenotypes.

Our study involves developing a statistical approach centered around gene pathways. This method incorporates estimations of Amyloid-β and tau tangle brain levels, along with memory scores from the MMSE, and integrates them into a comprehensive statistical model. Additionally, we include disease-relevant traits such as age, sex, education, and APOE genotype in this model. The primary goal is to rank the pathways that undergo alterations in AD, considering the influence of each of the biomarkers. By doing so, we can effectively identify the shared pathways across the three domains: amyloid, tau, and MMSE scores.

Moreover, our approach factors in the unique characteristics of human subjects, including age-specific, sex-specific, education-specific, and APOE genotype-specific differences. It also evaluates the significance of the relationship between pathway-level interactions and the presence of AD in relation to each of these factors. We conduct analyses using human transcriptomic data on Alzheimer’s disease progression and explore interactions with the APOE genotype.

To identify relevant gene sets [12] introduced Gene Set Enrichment Analysis (GSEA), a robust analytical method for interpreting gene expression data from genome-wide RNA analysis. GSEA focuses on gene sets—groups of genes with shared biological function, chromosomal location, or regulation—and demonstrates effectiveness in identifying common biological pathways e.g. for cancer-related data sets, such as leukemia and lung cancer, where single-gene analysis falls short. This method using the Kolmogorov-Smirnov statistics has a limitation in that it does not account for gene-gene interactions. Still, it has spurred the development of many other methods [13]. Other statistics have been proposed as well, e.g. the gene set likelihood ratio test (gsLRT), which uses a logistic regression model [14]. However, this particular model has its limitations as it is only applicable to binary outcome variables. In contrast, our study deals with continuous-scale variables, specifically amyloid, tau, and MMSE scores. As a result, we require a different model that can accommodate such continuous data. Additionally, we incorporated transcriptomic data, encompassing more than 20 thousand genes, as opposed to the mere 142 genes used in the referenced gsLRT study involving proteomics.

## Methodology

Our proposition introduces a novel approach utilizing all three phenotypes and biomarkers (amyloid, tau, and MMSE scores) to identify shared and meaningful pathways associated with them. Our methodology harnesses gene-level data, encompassing transcriptomics. By utilizing this data, we generate significance scores at the biological pathway level, serving as indicators of the novelty of observed experimental outcomes.

### Data and preprocessing

The data sample was taken from a subset of the Religious Orders Study and Rush Memory and Aging Project (ROSMAP) dataset [15–17] that had RNAseq data available from the dorsolateral frontal cortex. ROS has enlisted nuns and brothers since 1994. MAP recruited individuals from the northern Illinois region since 1997. Both studies were run by the same investigators using similar data collection techniques. Thus, the results from both were comparable. For the analyses reported in this paper, the clinical consensus diagnoses of Alzheimer’s disease or mild cognitive impairment were used to define a case while the diagnosis of no cognitive impairment/no impaired domains defined controls. Additional covariates for the statistical models were age, sex, education, and APOE genotype. The total sample with both gene expression and clinical data contained 634 subjects, with 433 cases and 201 controls. Demographic information for the sample is summarized in Table 1.

**Table 1.**
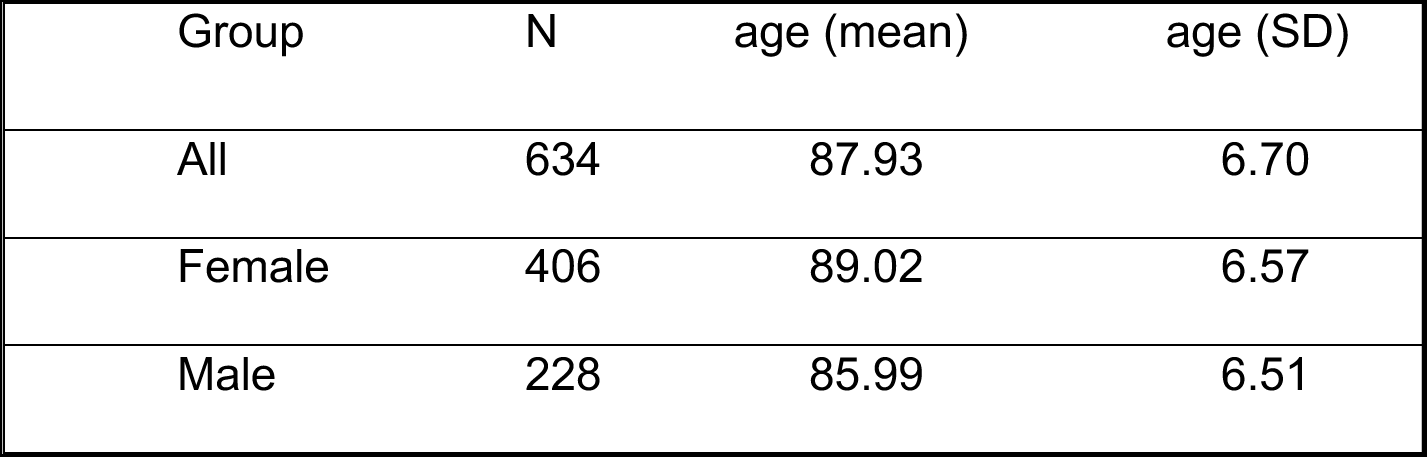
Demographic Information of Participants by Gender. This table presents the demographic information for a sample of 634 participants, categorized by gender. It includes the mean age and standard deviation (SD) for each group.

| Group | N | age (mean) | age (SD) |
| --- | --- | --- | --- |
| All | 634 | 87.93 | 6.70 |
| Female | 406 | 89.02 | 6.57 |
| Male | 228 | 85.99 | 6.51 |

In the context of [18], the amyloid and tangles definitions and measurements are as follows.

The overall amyloid level is determined as the mean percentage of cortex occupied by Amyloid-β protein in eight specific brain regions. This measurement is obtained through molecular-specific immunohistochemistry, where the Amyloid-β protein is targeted and quantified using image analysis techniques. The Amyloid-β score is calculated in eight brain regions, namely the hippocampus, entorhinal cortex, midfrontal cortex, inferior temporal cortex, angular gyrus, calcarine cortex, anterior cingulate cortex, and superior frontal cortex. At least four of these regions are required to calculate the mean Amyloid-βscore.

Tangle density is determined as the mean density of neuronal neurofibrillary tangles in eight specific brain regions. These tangles are identified using molecular-specific immunohistochemistry, employing antibodies specific to abnormally phosphorylated Tau protein, known as AT8. The cortical density of tangles is measured per square millimeter using systematic sampling. The tangle score is calculated as the mean density in eight brain regions, including the hippocampus, entorhinal cortex, midfrontal cortex, inferior temporal cortex, angular gyrus, calcarine cortex, anterior cingulate cortex, and superior frontal cortex. A minimum of four regions is necessary to compute the mean tangle density.

Clinical phenotypic information for the sample is summarized in Table 2.

**Table 2.** Clinical Phenotypic Data for Different Cognitive States. This table presents key clinical phenotypic data for different cognitive states in a diverse sample population. The study includes individuals with no cognitive impairment, mild cognitive impairment (MCI), Alzheimer’s dementia (AD), and other primary causes of dementia. The table provides mean scores and standard deviations for Mini-Mental State Examination (MMSE), amyloid levels, and log(tangles) for each cognitive state/phenotype. The findings highlight distinct cognitive profiles and potential biomarkers associated with various cognitive conditions, contributing to better understanding and targeted interventions for cognitive disorders.

| Cognitive state/Phenotypes | N | MMSE | MMSE | Amyloid | Amyloid | log(tangles) | log(tangles) |
| --- | --- | --- | --- | --- | --- | --- | --- |
|  | Rows | (mean) | (SD) | (mean) | (SD) | (mean) | (SD) |
| No cognitive impairment | 201 | 28.21 | 1.6 | 2.78 | 3.44 | 0.32 | 1.58 |
| MCI: Mild cognitive impairment (one impaired domain) and NO other cause of CI | 158 | 25.89 | 3.38 | 4.03 | 4.63 | 1.07 | 1.28 |
| MCI: Mild cognitive impairment (One impaired domain) AND another cause of CI | 10 | 25.19 | 3.73 | 1.35 | 1.33 | -0.03 | 1.3 |
| AD: Alzheimer's dementia and NO other cause of CI (NINCDS PROB AD) | 220 | 13.49 | 8.22 | 5.72 | 4.36 | 1.83 | 1.25 |
| AD: Alzheimer's dementia AND another cause of CI (NINCDS POSS AD) | 33 | 15.49 | 7.94 | 3.62 | 3.69 | 1.41 | 0.97 |
| Other dementia: Other primary cause of dementia | 12 | 15.17 | 5.47 | 2.59 | 3.6 | 0.5 | 1.31 |

The RNAseq data was obtained from the Accelerating Medicines Partnership Program for Alzheimer’s Disease Data Knowledge Portal (https://adknowledgeportal.synapse.org/), specifically, the RNA-seq Harmonization study (https://www.synapse.org/#!Synapse:syn9702085). The ROSMAP data from this study was used to create a combined dataset of RNA-seq data in combination with the three clinical phenotypes of MMSE, amyloid burden, and tangles [19]. The RNA-seq Harmonization study has the goal of creating an RNA-seq database based on a consensus set of analytical tools. The methodological details of the RNA-seq processing are given in Wan et al [20] and at the RNA-Seq reprocessing study website for the ROSMAP project (https://www.synapse.org/#!Synapse:syn8456629). In brief, RNA was extracted from samples consisting of approximately 100 mg of gray matter tissue from the dorsolateral prefrontal cortex. The RNA samples were prepared and sequenced as described in [19]. The reprocessing of the RNAseq data was done using a consensus set of tools with only library type-specific parameters varying between pipelines. Picard (https://broadinstitute.github.io/picard/) was used to generate FASTQs from source BAMs.

Generated FASTQ reads were aligned to the GENCODE24 (GRCh38) reference genome using STAR [21] and gene counts were computed for each sample. To evaluate the quality of individual samples and to identify potentially important covariates for expression modeling, two sets of metrics were computed using the CollectAlignmentSummaryMetrics and CollectRnaSeqMetrics functions in Picard. To account for differences between samples, studies, experimental batch effects, and unwanted RNA-Seq specific technical variations library normalization and covariate adjustments for each study separately using fixed/mixed effects modeling. The workflow consists of the following steps: (i) gene filtering: Genes that are expressed more than 1 CPM (read Counts Per Million total reads) in at least 50% of samples in each tissue and diagnosis category were used for further analysis, (ii) conditional quantile normalization was applied to account for variations in gene length and GC content, (iii) sample outlier detection using principal component analysis and clustering, (iv) Covariates identification and adjustment, where confidence of sampling abundance were estimated using a weighted linear model using voom-limma package in Bioconductor [22]. For the differential expression analysis, fixed/mixed effect linear regression was used with the following models: gene expression ∼Diagnosis + Sex + covariates + (1| Donor) or gene expression ∼Diagnosis x Sex + covariates + (1|Donor), where each gene is linearly regressed independently with Diagnosis, a variable explaining the AD status of an individual, identified covariates, and donor information as a random effect. Observation weights (if any) were calculated using the voom-limma [22] pipeline such that observations with higher presumed precision will be up-weighted in the linear model fitting process.

The data used for the gsLRT analysis included the phenotypes MMSE, amyloid burden and log(tangles). For gene expression, normalized and filtered counts were used as per the description on the RNA-Seq reprocessing study webite (https://www.synapse.org/#!Synapse:syn8456629). The specific datasets used for this study are: RNAseq data is obtained from (https://www.synapse.org/#!Synapse:syn8456638), covariates are recorded in (https://www.synapse.org/#!Synapse:syn11024258), differential expression results are from (https://www.synapse.org/#!Synapse:syn8456721) and clinical phenotypic data was obtained from the ROSMAP data sharing resource (https://www.radc.rush.edu/). These datasets were organized into the specific matrices for gene expression results, covariates, and phenotypes required by the gsLRT program.

### Data and Code Availability

ROSMAP resources can be requested at https://www.radc.rush.edu. The Code is available at https://github.com/Ali-Mahzarnia/gslrt2.

### Statistical Model

To derive the pathways associated with each AD related phenotypes (amyloid, tau, or MMSE scores), we define the following hypothesis tests. In our models, we utilize a matrix 𝐺 ∈ ℝ*^n^*^×*m*^ to represent measurement values (e.g., RNA expression) for m genes and n samples. Additionally, we employ a matrix, 𝑋 ∈ ℝ*^n^*^×*d*^, which contains d=4 covariates such as "sex," years of education”, “APOE Genotype” (APOE2,3,4), and “age at death” associated with the samples. Moreover, we utilize a continuous vector 𝑦 ∈ ℝ*^n^*^×1^ for the n samples that is the phenotype measurements (amyloid, tau, or MMSE scores). To conduct data analysis, we employ nested models for each gene 𝑔*_j_*, 𝑗 = 1, . . ., 𝑚 and all samples 𝑖 = 1, . . ., 𝑛 as outlined below:

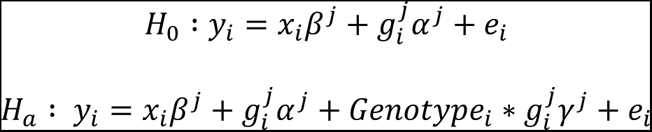

where 𝑒*_i_*∼ 𝑁(0, 𝜎). We denote the maximum likelihood estimator of the two above models with 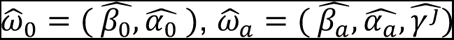.

In order to assess the enhanced explanatory capacity of the interaction term between the gene measurement profile (𝑔*_j_*) and genotype (APOE2,3,4), in contrast to the simpler model that merely includes the covariate matrix X and (𝑔*_j_*), we introduce statistical measures that evaluate the disparity in the models via log-likelihoods at the gene level. These statistics act as our metric for quantifying the supplementary explanatory power.

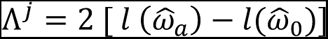

where 𝑙(.), is the log-likelihood. We define the enrichment score for gene set 𝐺*_j_*:

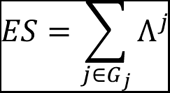

We calculate p-values for enrichment scores through nperm = 10,000 permutation sampling, which provides more conservative estimates but is computationally intensive. Since gene set tests share overlapping membership and exhibit interdependence, the presented p-values in this context and subsequent tables have not been adjusted for multiple testing. Consequently, they do not possess theoretical guarantees for controlling the False Discovery Rate (FDR). Once the pathways are sorted by their significance or ES, identifying the shared pathways among the three model runs becomes possible.

The model utilized in this study closely resembles that developed by Bryan et al [14], based on a logistic model, whereas our model accommodates a continuous outcome variable.

## Results

Using public resources from RNA-seq analyses of prefrontal cortex in ROSMAP participants, we have identified pathways associated for Amyloid-β, tau tangles and MMSE, accounting for covariates i.e. sex, genotype, education, age, and the interactions between RNA expression levels and APOE genotype.

We have identified significant pathways when examining the outcome of Amyloid-β. We have chosen a p-value threshold of 0.1 instead of the conventional 0.05 for hypothesis decision-making in order to capture a broader range of potentially significant pathways. This slightly relaxed threshold allows for a more inclusive analysis, potentially uncovering additional pathways that may contribute to our understanding of the relationship between Amyloid-β and the identified pathways. Table 3 and Figure 1 present the 14 significant pathways associated with Amyloid-β, such as TRANSLOCASE ACTIVITY, C3HC4 TYPE RING FINGER DOMAIN BINDING, INSULIN RECEPTOR BINDING, and VASCULAR ENDOTHELIAL GROWTH FACTOR RECEPTOR 2 BINDING.

**Figure 1.**
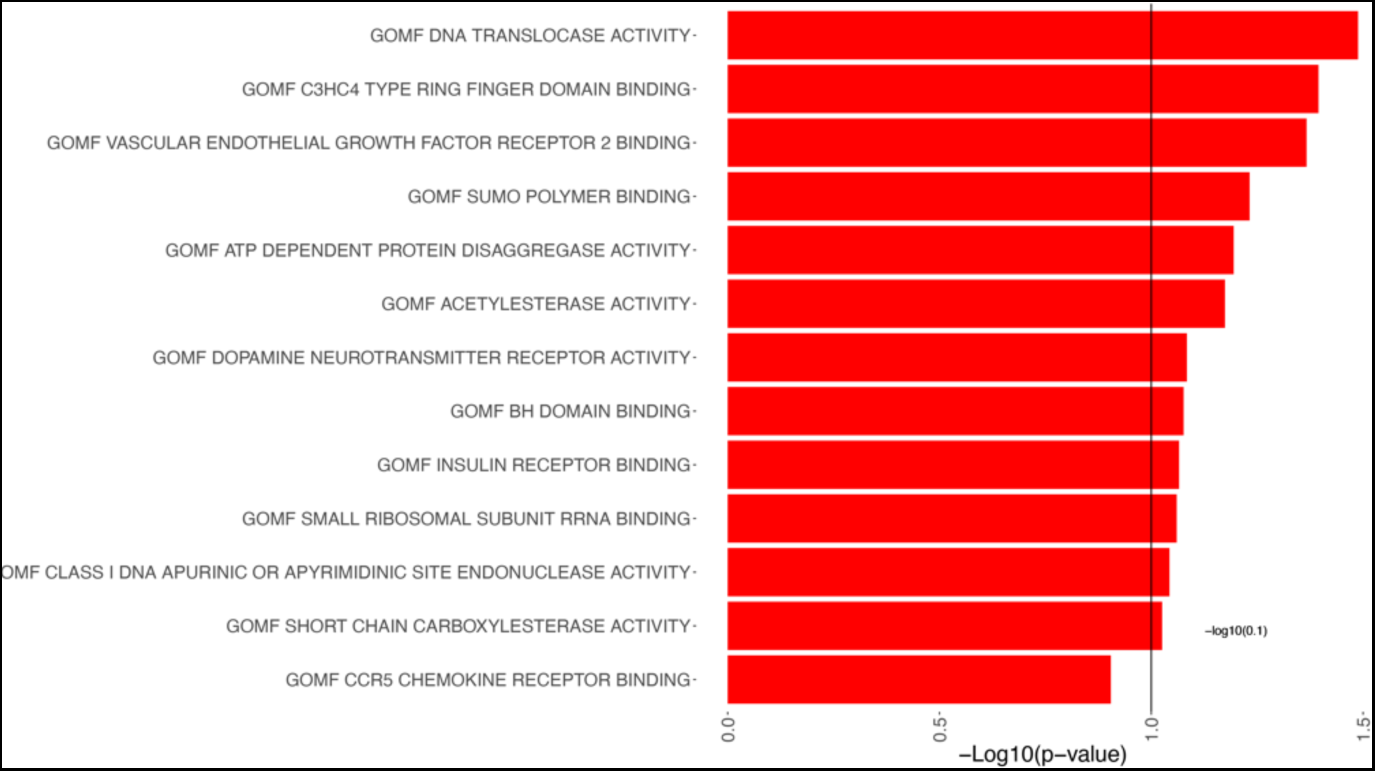
Pathway Analysis of Amyloid-β. Each pathway is represented on the y-axis, while the corresponding -log10(P-value) is represented by bars parallel to the x-axis, positioned in front of each pathway. The length of the bars reflects the statistical significance of the pathways with amyloid burden; longer bars indicating greater significance. This visualization allows for a quick assessment of the significance levels for each pathway, aiding in the identification of key pathways associated with Amyloid-β.

**Table 3.** Significant Pathways Associated with Amyloid-β Using a P-value Threshold of 0.1. This table presents the pathways that show significance (p-values smaller than 0.1) in relation to Amyloid-β,. The pathways listed provide insights into potential biological mechanisms and molecular processes associated with Amyloid-β.

| Gene Pathways | p-value | ES |
| --- | --- | --- |
| GOMF DNA TRANSLOCASE ACTIVITY | 0.032 | 3.761 |
| GOMF C3HC4 TYPE RING FINGER DOMAIN BINDING | 0.040 | 3.047 |
| GOMF VASCULAR ENDOTHELIAL GROWTH FACTOR RECEPTOR 2 BINDING | 0.043 | 2.445 |
| GOMF SUMO POLYMER BINDING | 0.059 | 2.637 |
| GOMF ATP DEPENDENT PROTEIN DISAGGREGASE ACTIVITY | 0.064 | 2.468 |
| GOMF ACETYLESTERASE ACTIVITY | 0.067 | 2.315 |
| GOMF DOPAMINE NEUROTRANSMITTER RECEPTOR ACTIVITY | 0.082 | 2.383 |
| GOMF BH DOMAIN BINDING | 0.084 | 2.074 |
| GOMF INSULIN RECEPTOR BINDING | 0.086 | 2.217 |
| GOMF SMALL RIBOSOMAL SUBUNIT RRNA BINDING | 0.087 | 2.002 |
| GOMF CLASS I DNA APURINIC OR APYRIMIDINIC SITE ENDONUCLEASE ACTIVITY | 0.091 | 2.298 |
| GOMF SHORT CHAIN CARBOXYLESTERASE ACTIVITY | 0.094 | 2.001 |
| GOMF VASCULAR ENDOTHELIAL GROWTH FACTOR RECEPTOR BINDING | 0.098 | 1.787 |
| GOMF ALPHA N ACETYL GALACTOSAMINIDE ALPHA 2 6 SIALYLTRANSFERASE ACTIVITY | 0.099 | 1.869 |

The 5 significant pathways associated with log(tangles) are presented in Table 4 and illustrated in Figure 2 such as BITTER TASTE RECEPTOR ACTIVITY, PROTEIN GLUTAMINE GAMMA GLUTAMYLTRANSFERASE ACTIVITY, TASTE RECEPTOR ACTIVITY, and VASCULAR ENDOTHELIAL GROWTH FACTOR RECEPTOR 2 BINDING.

**Figure 2.**
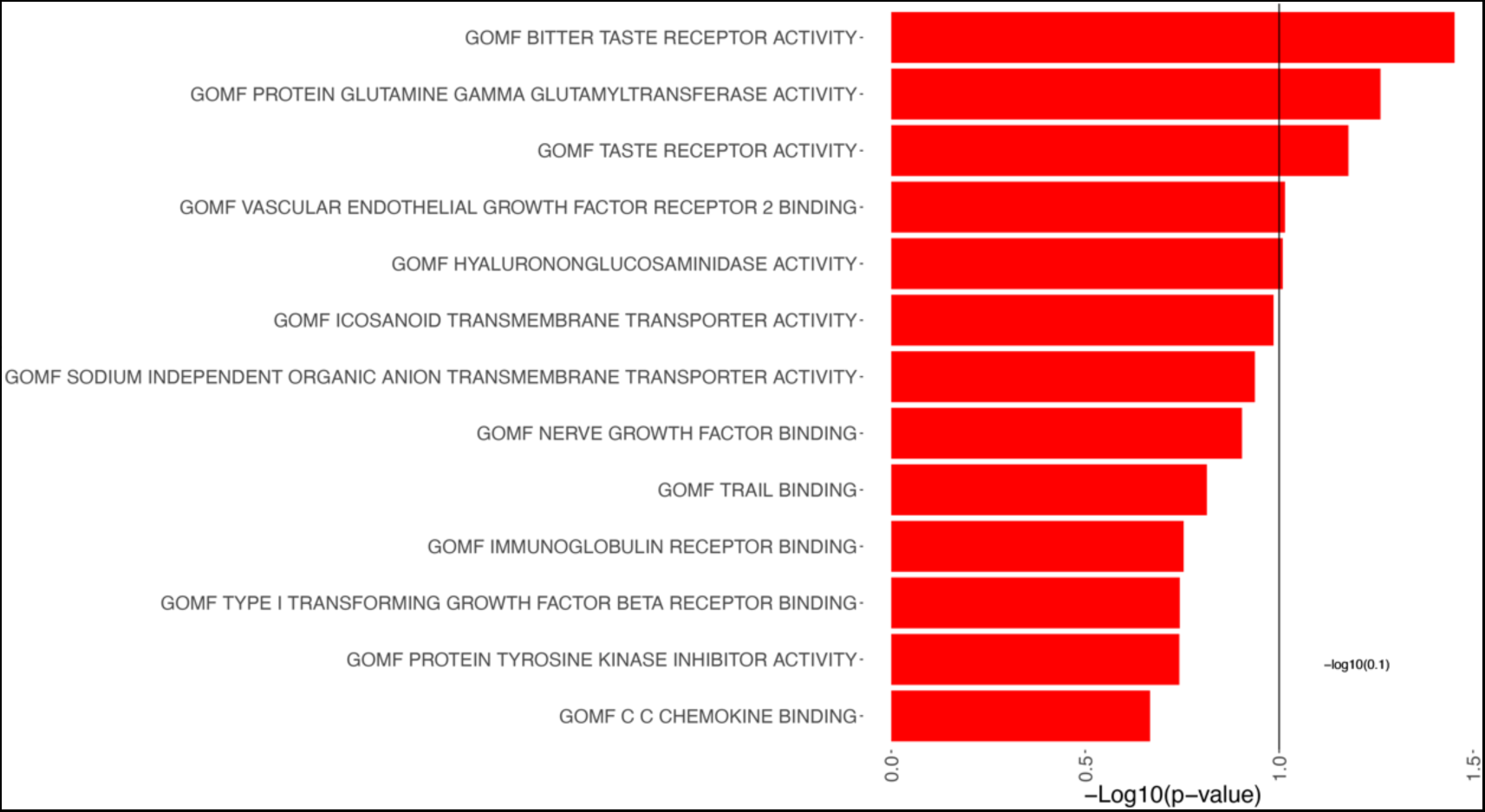
Pathway Analysis of log(tangles). This figure presents the results of the pathway significance analysis for log(tangles). Each pathway is displayed on the y-axis, while bars parallel to the x-axis represent the corresponding -log10(P-value). The length of each bar reflects the statistical significance of the pathway association with log(tangles). Longer bars indicate greater significance/smaller p values.

**Table 4.** Significant Pathways Associated with log(tangles). This table shows the pathways that exhibited statistical significance (p-values smaller than 0.1) in relation to log(tangles).

| Gene Pathways | p-value | ES |
| --- | --- | --- |
| GOMF BITTER TASTE RECEPTOR ACTIVITY | 0.035 | 3.547 |
| GOMF PROTEIN GLUTAMINE GAMMA GLUTAMYLTRANSFERASE ACTIVITY | 0.055 | 2.658 |
| GOMF TASTE RECEPTOR ACTIVITY | 0.066 | 2.486 |
| GOMF VASCULAR ENDOTHELIAL GROWTH FACTOR RECEPTOR 2 BINDING | 0.097 | 1.910 |
| GOMF HYALURONONGLUCOSAMINIDASE ACTIVITY | 0.098 | 2.074 |

In addition to the pathways depicted in Figure 3, Table 5 includes the top candidate of such pathways, which represent a subset of significant associations (with p-values smaller than 0.1) with the Mini-Mental State Examination (MMSE) outcome. These results highlight specific pathways that exhibit statistical significance in association with MMSE. Figure 3 provides a partial representation of 174 top pathways, including TRAIL BINDING, VASCULAR ENDOTHELIAL GROWTH FACTOR RECEPTOR BINDING, VASCULAR ENDOTHELIAL GROWTH FACTOR RECEPTOR 2 BINDING

**Figure 3.**
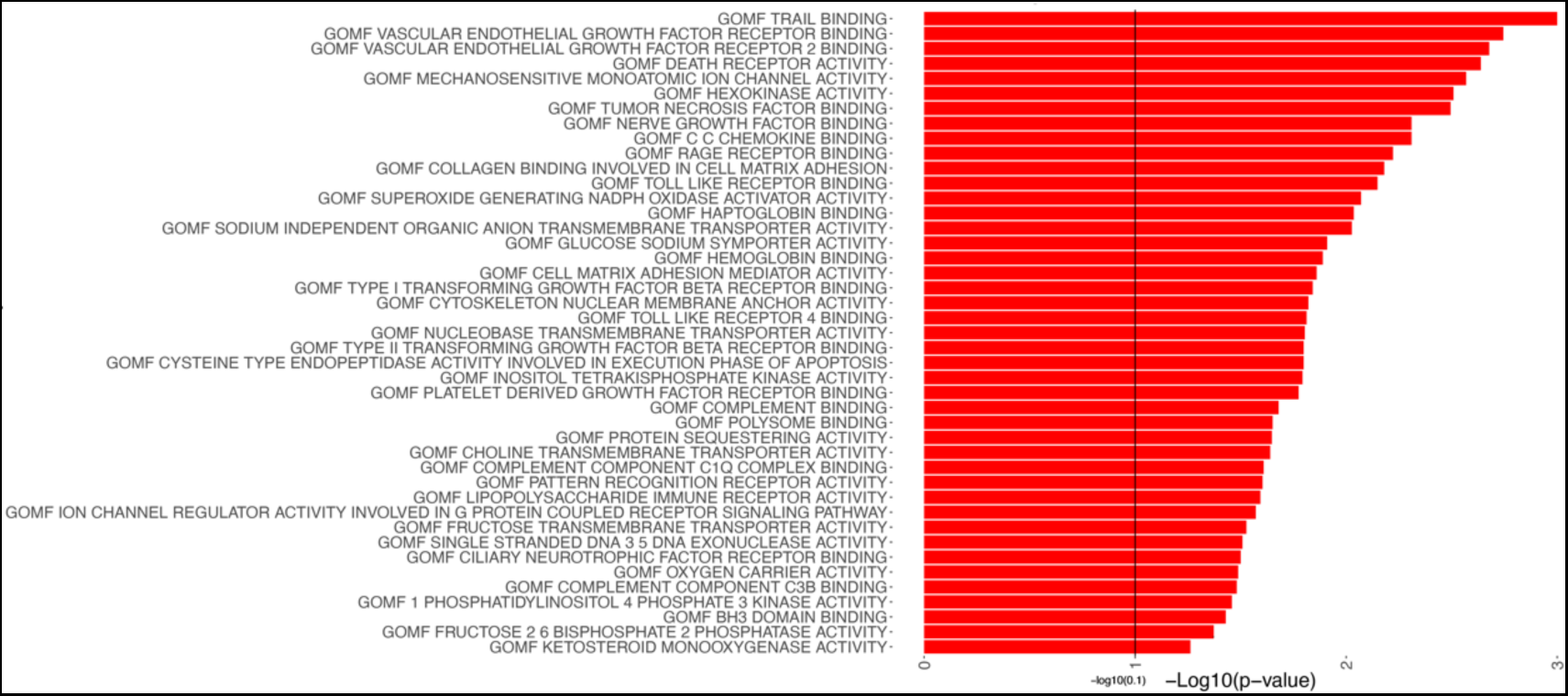
Pathway Analysis of MMSE. This figure shows a portion of the pathway significance analysis results for MMSE. Each pathway is plotted on the y-axis, accompanied by bars parallel to the x-axis that represent the corresponding -log10(P-value). The length of each bar reflects the statistical significance of the pathway, with longer bars indicating higher significance. This visual representation enables a quick evaluation of the significance levels associated with each pathway. It assists in identifying key pathways that are linked to MMSE, contributing to a better understanding of the underlying mechanisms influencing MMSE.

**Table 5.**
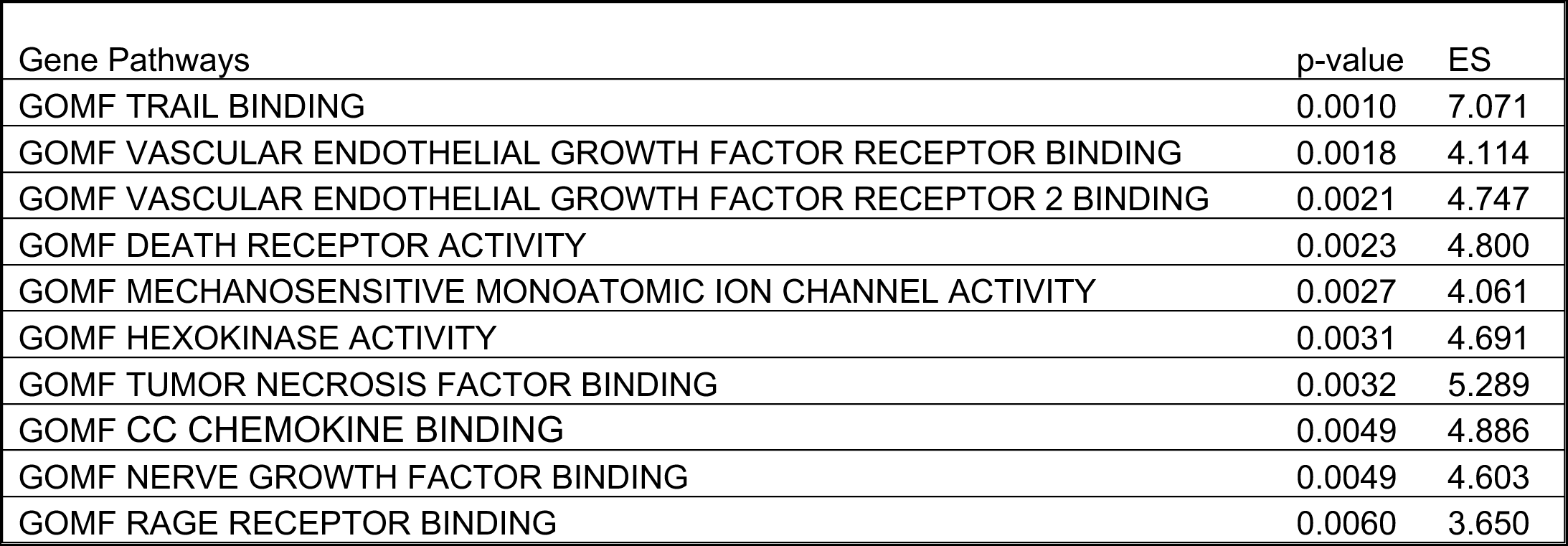
Significant Gene Pathways Associated with MMSE Outcome. It highlights the top candidate pathways, including TRAIL BINDING and VASCULAR ENDOTHELIAL GROWTH FACTOR RECEPTOR BINDING, among others.

Finally, we investigated the shared pathways among all 3 studies involving Amyloid-β, log(tangles), and MMSE, shown in Figure 4. This analysis allows us to gain insights into the shared biological mechanisms and molecular processes that may contribute to the interplay between Amyloid-β, log(tangles), and MMSE.

**Figure 4.**
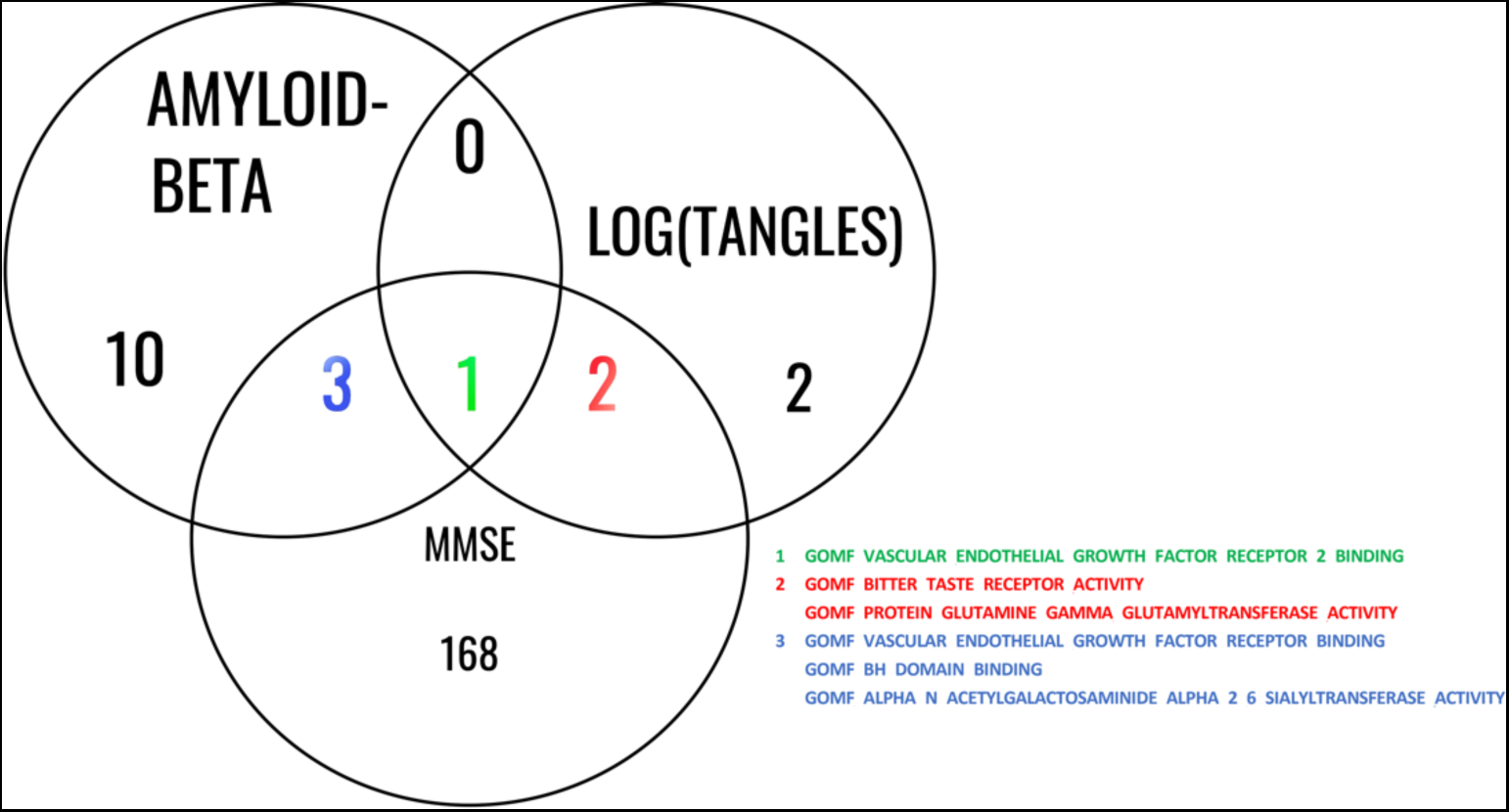
Common Significant Pathways Between for Amyloid-β, log(tangles), and MMSE. This Venn diagram illustrates the common pathways identified among the studies involving Amyloid-β, log(tangles), and MMSE. The diagram consists of overlapping circles that represent each study, with labeled sections indicating the shared pathways among them. The shared pathways are listed within the diagram, providing a concise overview of the biological processes and molecular mechanisms that are consistently implicated across these phenotypes. This analysis highlights the interconnectedness of these factors and underlying pathways that contribute to the associations between Amyloid-β, log(tangles), and MMSE.

Results for the comparisons of expression differences between control and AD samples are shown in Table 6 as reported in the AMP-AD analysis of the dataset (https://www.synapse.org/#!Synapse:syn8456721). Higher expression in AD samples relative to normal cognition is observed for VEGFA, VEGFB, VEGFD, PGF and FLT1 while lower expression in AD samples relative to controls is observed for FLT4, KDR, NRP1 and NRP2. The only significant difference between the two groups after adjusting for multiple comparisons is for VEGFB.

**Table 6.** Differential Expression of VEGF family related Genes in Alzheimer’s Disease (AD) Samples Compared to Controls. This table presents the results of a comparative analysis between control and Alzheimer’s disease (AD) samples, focusing on the expression differences of VEGF-family--related genes.

| Gene | Comparison | logFC | CI.L | CI.R | AveExpr | t | P.Value | adj.P.Val |
| --- | --- | --- | --- | --- | --- | --- | --- | --- |
| VEGFA | AD-CONTROL | 0.16 | -0.08 | 0.40 | 1.86 | 1.28 | 0.20 | 0.43 |
| VEGFB | AD-CONTROL | 0.10 | 0.03 | 0.16 | 5.67 | 2.81 | 0.01 | 0.05 |
| VEGFD | AD-CONTROL | 0.02 | -0.07 | 0.12 | 1.10 | 0.50 | 0.62 | 0.79 |
| PGF | AD-CONTROL | 0.15 | -0.02 | 0.32 | 0.17 | 1.69 | 0.09 | 0.26 |
| FLT1 | AD-CONTROL | 0.14 | -0.05 | 0.32 | 3.86 | 1.44 | 0.15 | 0.36 |
| FLT4 | AD-CONTROL | -0.03 | -0.15 | 0.09 | 0.15 | -0.54 | 0.59 | 0.78 |
| KDR | AD-CONTROL | -0.07 | -0.20 | 0.07 | 1.05 | -1.00 | 0.32 | 0.55 |
| NRP1 | AD-CONTROL | -0.07 | -0.18 | 0.03 | 2.14 | -1.41 | 0.16 | 0.37 |
| NRP2 | AD-CONTROL | -0.02 | -0.10 | 0.06 | 1.37 | -0.58 | 0.56 | 0.75 |

## Discussion

The progression of Alzheimer’s disease (AD) manifests through changes in biomarkers that reflect abnormal protein expression, such as Amyloid-β, phosphorylated tau, as well as clinically measurable symptoms including memory decline. In our study, we developed new methods to reveal pathways related to changes in brain RNA-Seq for each of these two neuropathological biomarkers, as well as for MMSE. Surprisingly we observed a larger number of significant pathways for gene expression association with MMSE (174 pathways) than those associated with the two hallmark biomarkers for AD, Amyloid-β (14 pathways), and tau tangles (5 pathways). More importantly, we identified a single pathway, vascular endothelial grown factor receptor binding (VEGF-RB) that was associated with differences reflective of all three phenotypes analyzed by the continuous gene set likelihood ratio test (gsLRT).

The methodological advance of this study was the extension of gsLRT from binary (e.g. disease/control status) to continuous value phenotypes. Our study extends prior work by analyzing three continuous scale phenotypes (Amyloid-β, tau tangles, MMSE) in a pathway analysis that accounts for the covariates age, sex and APOE genotype (gsLRT for continuous phenotypes).

Among the pathways identified as significant for Amyloid-β we noted several candidate pathways that support changes in DNA repair ability (GOMF DNA TRANSLOCASE ACTIVITY) [23], cell mediated immunity (GOMF C3HC4 TYPE RING FINGER DOMAIN BINDING) [24], apoptosis (GOMF BH DOMAIN BINDING) [25], protein synthesis (GOMF SMALL RIBOSOMAL SUBUNIT RRNA BINDING) [26] and disaggregation (GOMF ATP DEPENDENT PROTEIN DISAGGREGASE ACTIVITY), as well as insulin signaling (GOMF INSULIN RECEPTOR BINDING) [27, 28], which have all been connected to AD.

Among the pathways identified as significant for tau tangles we noted two that support alterations in sensory processing, pointing to taste (GOMF BITTER TASTE RECEPTOR ACTIVITY; GOMF TASTE RECEPTOR ACTIVITY). Recent studies have shown alterations in the sour taste [29], while here we identified changes in pathways associated with bitter taste, and taste in general. Since taste and olfaction are closely linked, these results suggest possible changes in olfactory function. Among sensory changes in AD, olfaction has been proposed as one of the more promising biomarkers for early detection [30]. We also noted pathways pointing to differences in the transfer of amino acids across the membrane, cell survival during oxidative stress (GOMF PROTEIN GLUTAMINE GAMMA GLUTAMYLTRANSFERASE ACTIVITY) [31], and glutathione homeostasis, relevant to several neurodegenerative diseases, such as AD, PD and ALS [31, 32]. Finally, the role of extracellular matrix was suggested by the presence of the GOMF HYALURONONGLUCOSAMINIDASE ACTIVITY pathway, which has a less understood but complex role in aging and disease [33, 34].

Among the large number of pathways related to MMSE, we noted top candidates involved in apoptosis through TRAIL which binds to death receptors, suggesting a relation with immune related mechanisms [35].

Amyloid-β and tau biomarkers in combination with measures of neurodegeneration have been associated with the progression of AD neuropathology and memory loss [36–41]. We note that neuropsychological measures such as MMSE can reflect other causes of cognitive impairment including damage to the cerebrovascular system although individuals with moderate to severe Alzheimer’s disease tend to have MMSE scores less than 15 [42].

The role of the VEGF signaling family in neurodegeneration and Alzheimer’s disease has been extensively studied including with multiomic approaches that analyzed bulk RNA sequencing data, single nucleus sequencing data, and mass spectrometry proteomics data [10]. As a consequence of the numerous pathways that intersect with VEGF receptor signaling, it has been difficult to identify the specific receptors and molecules that associate with disease endophenotypes or covariates including age and sex.

VEGF includes a family of five ligands (VEGFA, VEGFB, VEGFC, VEGFD, and PGF), three tyrosine kinase receptors (FLT1, FLT4, and KDR), and two modulating receptors (NRP1 and NRP2). The pathway identified as significantly associated with the three phenotypes (GOMF VASCULAR ENDOTHELIAL GROWTH FACTOR RECEPTOR BINDING (GO:0005172) includes the ligands and several other genes. The pathway GOMF VASCULAR ENDOTHELIAL GROWTH FACTOR RECEPTOR ACTIVITY (GO:0005021) includes the tyrosine kinase receptors and modulating receptors).

The presence of VEGF alongside Amyloid-β plaques in AD brains and its strong binding to Amyloid-β suggest that VEGF may contribute to neurodegeneration and vascular dysfunction [43]. Additionally, Amyloid-β inhibits VEGF receptor signaling, impairing angiogenesis [44]. VEGF accumulation around amyloid plaques interacts directly with Amyloid-β, rescuing synaptic dysfunction caused by the toxic Amyloid-β oligomers [45].

VEGF also interacted with tau and amyloid-β42, predicting hippocampal atrophy and memory decline. Another study [46] revealed that VEGF genes, particularly FLT4 and FLT1, were associated with AD neuropathology and cognition. Higher levels of VEGF were associated with slower hippocampal atrophy and better cognitive function [47]. These findings emphasize the importance of understanding the relationship between VEGF, tau pathology, and AD for potential therapeutic interventions.

Several studies have investigated the role of vascular endothelial growth factor (VEGF) in cognitive impairment. For example one study found that VEGF AA genotype is associated with an increased risk of developing AD and MCI, while higher VEGF levels are observed in AD patients [48]. Another study found lower VEGF levels in AD patients and amnestic MCI patients compared to controls, correlating with cognitive decline [49]. Additionally, [50] finds that higher serum VEGF levels in ischemic stroke patients are associated with post-stroke cognitive impairment. Conversely, [51] shows that VEGF signaling is crucial for maintaining cognition and neurogenesis, cautioning against inhibiting VEGF signaling. Interestingly, [52] demonstrates that VEGF levels increase during the early stage of AD but decrease as the disease progresses, suggesting a link between VEGF levels and cognitive decline. VEGF produced by macrophages plays a role in preserving cognitive function in obesity, which can be a risk factor during aging and AD [53] However the literature is still controversial and more work is needed to understand the role of various VEGF isoforms role in modulating cognition [54] .

The ensemble of molecules in the VEGF pathways and their interactions have been reported to have varied effects on AD phenotypes. The members of the VEGF measured in the brain and blood have been characterized with respect to cognitive performance, neural and cerebrovascular pathology, and CSF biomarkers [11, 55]. Blood and brain VEGFA has been reported to be protective against memory impairment and brain atrophy in AD [10]. Higher expression levels of VEGFB, PGF, FLT1 and FLT4 are reported to be associated with faster cognitive decline and greater neuropathological lesion development [10, 55] and our data supported this direction of difference in expression levels. The VEGF family is involved in multiple signaling pathways, leading to potentially different effects on AD-related phenotypes; for example, VEGFA can signal through KDR or FLT1 where the receptors can elicit effects in opposing directions [10]. In a study of microglial control of astrocytes in response to microbial metabolites, microglial VEGFB was shown to trigger FLT1 signaling in astrocytes and promote CNS inflammation [56]. Neutropilin expression (NRP1 and NRP2) was decreased AD samples relative to samples from individuals with normal cognition in agreement with prior evidence [10]. NRP1 and NRP2 have well-established roles in angiogenesis[10, 57]. Moreover, interactions between VEGF family proteins with APOE have also been reported. Higher levels of VEGFA were reported to be associated with worse outcomes among APOE E4 carriers and better outcomes among non-E4 carriers [10].

Our study has several strengths. First, the extension of the gsLRT to continuous phenotypes allowed a higher statistical level (continuous vs. nominal) of data for the pathway analysis. The relatively large sample size where data was available for all three phenotypes allowed for comparisons at the gene and pathway level. For the pathway/signature analysis, well-established databases including gene ontology were used to enable replication studies and other future work.

A limitation of the analysis is that the phenotypes are measured at a single time point. Longitudinal data for the Amyloid-β and tau biomarkers are not available from post-mortem brains, though nuclear imaging or fluid biomarkers present a great promise for the future of such longitudinal studies. MMSE is measured over time, so another possible phenotype to consider would be the decline in MMSE from baseline to death. Studies that examine changes in CSF and blood biomarkers for VEGF over time will provide information on temporal relationships between VEGF family mRNA and protein concentrations and biomarker changes and can open new avenues for exploiting its therapeutic potential [10].

Our study proposes a method for pathway identification using continuous phenotypes and public data bases on RNA-Seq, but can potentially be used for proteomic analyses, or extended to multinetwork omic studies and make use of extensive public data base resources or de novo analyses to better understand the mechanistic substrates for neurodegenerative diseases.

## Acknowledgment

We thank all the participants of ROS and MAP studies. These studies were funded by the National Institute on Aging: P30AG010161 ADCC, P30AG072975 ADRC, R01AG015819 RISK, R01AG017917 MAP, U01AG46152 AMP-AD Pipeline I, U01AG61356 AMP-AD Pipeline II. Furthermore, we are grateful to Dr. Jordan Bryan for helpful discussions.

